# Metatranscriptomic analysis reveals synergistic activities of comammox and anammox bacteria in full-scale attached growth nitrogen removal system

**DOI:** 10.1101/2024.01.08.574720

**Authors:** Juliet Johnston, Katherine Vilardi, Irmarie Cotto, Ashwin Sudarshan, Kaiqin Bian, Stephanie Klaus, Megan Bachmann, Mike Parsons, Christopher Wilson, Charles Bott, Ameet Pinto

## Abstract

0-

Leveraging comammox *Nitrospira* and anammox bacteria for shortcut nitrogen removal can drastically lower the carbon footprint of wastewater treatment facilities by decreasing aeration energy, carbon, alkalinity, and tank volume requirements while also potentially reducing nitrous oxide emissions. However, their co-occurrence as dominant nitrifying bacteria is rarely reported in full-scale wastewater treatment. As a result, there is poor understanding of how operational parameters, in particular dissolved oxygen, impact their activity and synergistic behavior. Here, we report the impact of dissolved oxygen concentration (DO = 2, 4, 6 mg/L) on the microbial community’s transcriptomic expression in a full-scale integrated fixed film activated sludge (IFAS) municipal wastewater treatment facility predominantly performed by comammox *Nitrospira* and anammox bacterial populations. 16S rRNA transcript compositions revealed anammox bacteria and *Nitrospira* were significantly more active in IFAS biofilms compared to suspended sludge biomass. In IFAS biofilms, anammox bacteria significantly increased *hzo* expression at lower dissolved oxygen concentrations and this increase was highly correlated with the *amoA* expression levels of comammox bacteria. Interestingly, the genes involved in nitrite oxidation by comammox bacteria were significantly more upregulated relative to the genes involved in ammonia oxidation with decreasing dissolved oxygen concentrations. Ultimately, our findings suggest that comammox *Nitrospira* supply anammox bacteria with nitrite via ammonia oxidation and that this synergistic behavior is dependent on dissolved oxygen concentrations.

**Synopsis:** Comammox bacteria differentially regulate ammonia and nitrite oxidation in response to dissolved oxygen concentration suggesting dissolved oxygen dependence of their synergistic nitrogen removal with anammox bacteria in IFAS biofilms.

## 1- Introduction

Recent studies have suggested that co-operation between comammox bacteria (CMX) and anammox bacteria (AMX) can successfully result in nitrogen removal from mainstream wastewater in both laboratory^1–3^ and full-scale systems^4^. Such cooperation between CMX and AMX could result in nitrogen removal with reduced oxygen and carbon demand, lower carbon, alkalinity, and tank volume requirements, while also producing significantly reducing biotic production of nitrous oxide (N_2_O)^5,6^ which is a highly potent greenhouse gas (GHG). This contrasts with conventional nitrogen removal systems, where estimates suggest 1-1.5% of incoming nitrogen is lost as fugitive N_2_O emissions^7^. Thus, the cooperation of CMX and AMX for wastewater treatment has the potential to significantly decarbonize wastewater treatment.

While CMX have been found in soils^8–10^, drinking water systems^11–13^, and traditional activated sludge systems^14–17^, they are far more likely to be prefer biofilms in attached growth systems and systems with long solids retention time (SRT)^18^ due to their slow growth rates^19,20^. Similarly, AMX may also prefer biofilms and granules due to their slower growth rates and because such aggregate environments may offer protection from their sensitivity to nitrite inhibition in nitrogen-rich wastewater sources^21,22^, while also preventing exposure to dissolved oxygen^23,24^. The cohabitation of CMX and AMX in biofilms provides an ideal scenario to drive nitrogen loss via their metabolic coupling. Specifically, CMX can drive down oxygen concentrations to ensure anaerobic environments for AMX while providing them with nitrite. CMX could further oxidize excessive nitrite to nitrate via nitrite oxidoreductase (*nxr*) activity, and potentially resupply nitrite from nitrate in anaerobic conditions since activity of the nitrite oxidoreductase enzyme is reversible. This sets up AMX to reduce ammonia and nitrite into dinitrogen gas establishing a strong symbiotic relationship.

Our previous work identified CMX and AMX are dominant nitrifying bacteria in biofilms in a full-scale integrated fixed film activated sludge (IFAS) process^4^; this was the first report to observe their potential cooperation in a full-scale municipal wastewater treatment system. In this study, we observed that anaerobic ammonia oxidation decreased, while aerobic ammonia oxidation to nitrate increased with increasing dissolved oxygen concentration. In the IFAS biofilms, two AMX bacterial populations associated with *Ca.* Brocadia, comprised 6.53 ± 0.34% of the population while CMX bacteria closely associated with *Ca. Nitrospira* nitrosa comprised 0.37 ± 0.03% of the population. This system still had an abundance of strict ammonia oxidizing bacteria (AOB) and nitrite oxidizing bacteria (NOB), particularly in the suspended phase making it unclear the extent to which organisms were likely supplying nitrite to AMX^18^.

The current study follows-up our previous metagenomic and kinetic investigation by systematically characterizing metatranscriptomic changes of the microbial communities in the suspended phase and IFAS biofilms in response to changing DO concentrations. To do this, we combine genome-centric metatranscriptomics as well as amplicon sequencing and quantitation of major functional gene expressions (*amoA* for AOB, *amoA* for CMX, *nxrB* for *Nitrospira* bacteria, and *hzo* for AMX) to determine (1) potential metabolic coupling between different nitrifying populations within this mixed community, and (2) transcriptomic responses of key nitrifying populations as a function of DO concentrations and growth mode (i.e., attached vs suspended phase). By understanding the activity of all nitrifying organisms (AOB, CMX, NOB, AMX) in a complex integrated fixed film activated sludge (IFAS) wastewater treatment facility, we can identify DO concentrations that produce the highest activity and potential synergy between CMX and AMX. Such insights have the potential to advance the development of mainstream partial-nitritation-anammox nitrogen removal systems with lower energy requirements and GHG emissions.

## 2- Materials and Methods

### 2.1 Overview of Experimental Design

A detailed overview of the wastewater treatment facility’s process flow diagrams, operational parameters, and conditions is reported in Vilardi et al 2023^4^. In brief, the secondary treatment consists of five-zones in series including an anaerobic stage followed by an anoxic stage receiving internal nitrate recycle divided into two cells, an aerobic stage with IFAS, and a small deaeration zone. Samples were taken at the anaerobic stage (R1), end of the anoxic zone (R3), and twice within the aerobic IFAS stage, at the beginning and the end (R4 and R5). Full-scale tank dissolved oxygen concentrations in the aerobic zone were stabilized for six-hours prior to sampling and experiments for multiple DO setpoints were run-on consecutive days within a week to eliminate potential confounding effects from changes in microbial community composition. Samples for DNA-based analyses were immediately stored in dry ice, while samples for RNA-based analyses were mixed at a 1:10 ratio of LifeGuard Soil Preservation (Qiagen, Hilden Germany) before being put on dry ice. At the end of each day’s sampling, all samples were stored at -80℃ until extraction.

### 2.2 Nucleic acid Extraction

DNA was extracted using protocols from the DNeasy PowerSoil Pro Kit with some modifications (Qiagen, Hilden, Germany) as described previously^4^. Extracted DNA was quantified using a Qubit 4 Fluorometer and standard protocols for dsDNA with High Sensitivity (ThermoFisher Scientific, Waltham, Massachusetts, USA). RNA was extracted using protocols from the Quick-RNA Fecal/Soil Microbe Microprep Kit with some modifications (Zymo, Irvine, California, USA). RNA samples were initially preserved with 1 mL of activated sludge (i.e., suspended phase) in 9 mL of LifeGuard Soil Preservation (Qiagen, Hilden Germany). This solution was thawed and centrifuged (16,000 g for 1-minute) to pellet the activated sludge. The 1 mL of activated sludge was then transferred into a 2 mL microcentrifuge tube and 1 mL of ultrapure DNAse/RNase free water was added. To remove residual RNA stabilization solutions, three rinses were performed where the samples were vortex for 30 seconds, centrifuged for 1-minute at 16,000 g, and 1.5 mL of supernatant was exchanged.

To remove the biomass from the IFAS plastic, IFAS and LifeGuard Soil Preservation were transferred into a 5 mL bead beating tube from DNeasy PowerWater kits for 5-minutes of vortexing (Qiagen, Hilden, Germany). Afterwards, 500 µL of biomass and LifeGuard Soil Preservation were transferred into a clean 2 mL microcentrifuge tube for rinsing to remove the RNA stabilization solution as described above.

Modifications made during the Quick-RNA Fecal/Soil Microbe Microprep Kit include using a FastPrep-24 homogenizer for 40 seconds at 6 m/s during bead beating. An additional DNAse_1 treatment was performed to degrade residual DNA (Zymo, Irvine, California, USA) followed by final elution of RNA in 50 µL of buffer. The eluted RNA and residual DNA were quantified via Qubit Fluorometer (ThermoFisher Scientific, Waltham, Massachusetts, USA). After quantification several aliquots were made where some eluted RNA was immediately frozen at - 80°C for metatranscriptomic sequencing, while another aliquot was processed for cDNA synthesis for RT-qPCR and amplicon sequencing before being frozen at -20°C.

### 2.4 Quantification of Transcripts and Genes

All samples were quantified for expression which included duplicate extractions from two technical replicates from all three dissolved oxygen settings, at all reactor locations (R1, R3, R4, and R5 for both suspended sludge biomass and IFAS). For DNA-based gene quantification, only technical duplicates from R1, R5 for suspended sludge biomass and IFAS were used as there was no significant change in community structure over the short experimental period (i.e., less than one week). The extracted RNA was synthesized into cDNA using standard protocols for SuperScript IV First-Strand Synthesis System for RT-PCR (Invitrogen, Waltham, Massachusetts, USA). The cDNA was quantified via Qubit Fluorometer (ThermoFisher Scientific, Waltham, Massachusetts, USA). RT-qPCR and qPCR were then performed on a QuantStudio 7 Flex (ThermoFisher Scientific, Waltham, Massachusetts, USA). The reaction mastermix contained 0.5 µL of both forward and reverse primers at 10 µM, 10 µL of Luna Universal qPCR Master Mix (New England Biolabs, Ipswich, Massachusetts, USA), 0.5 µL of template DNA/cDNA, and 8.5 µL of nuclease-free water. All primers were purchased by Integrated DNA Technologies (IDT, Coralville, Iowa, USA).

Primers and their thermocycling conditions are shown in **Supplemental Table 1** using standard protocols for 16S rRNA^25^, ammonia monooxygenase subunit A (*amoA*) for strict AOB^26^, *amoA* for *Ca.* Nitrospira nitrosa^27^, nitrite oxidoreductase subunit B (*nxrB*) for *Nitrospira* bacteria^28^, and hydrazine oxidoreductase (*hzo)* found in AMX^29^. Quantification data of transcripts is shown in **Supplemental Figure 1** with a comparison between reporting RT-qPCR results as log(copies/mL) and log(copies/g of Total Solids) in **Supplemental Figure 2.**

### 2.5 Amplicon Sequencing

Extracted DNA and cDNA were submitted to Georgia Institute of Technology’s Molecular Evolution Core where for sequencing using the Illumina MiSeq v3 kit PE300 using Illumina Nextera Adapter Sequences (Illumina Inc, San Diego, California, USA). The samples were demultiplexed and barcodes removed by the sequencing core. The raw sequences were then imported into RStudio v2023.06.2^30^ and processed using DADA2 v1.26^31^. In DADA2, 16S rRNA amplicons used standard protocols with a lower, default truncQ = 2. Other amplicons did not use any truncation, and during merging maxMismatch was increased to 5 to allow longer amplicons to have substantial overlap. Taxonomic assignments referenced the SILVA SSU 138.1 database^32^ and additionally checked in the with the MiDAS 5.0 database^33^. Merged reads outside ± 2 of the expected amplicon length were discarded. ASV’s were then consolidated into OTUs with a 97% identity cutoff using BioStrings v2.66.0^34^, dplyr v1.1.2^35^, tibble v3.2.1^36^, and DECIPHER v2.26.0^37^. Histograms of postprocessed amplicon reads are available in the supplemental materials (**Supplemental Figure 3**). Amplicon compositions for *amoA* for AOB, *amoA* for CMX, *nxrB* for *Nitrospira* sp., and *hzo* for AMX can be found in **Supplemental Figure 4** with top BLAST matches based on e-value with the reported sequence percent ID in similarity in **Supplemental Table 2.**

Absolute quantification of OTUs was estimated by multiplying the OTU relative abundance with RT-qPCR copies for each gene. The same primers were used for RT-qPCR amplification and amplicon sequencing to ensure parity with previous similar studies^38–42^.

### 2.6 Metatranscriptomic sequencing and data processing

A limited subset of samples were submitted for metatranscriptomic sequencing in duplicate, limited to only DO concentrations of 2 mg/L and 6 mg/L across reactor locations R1, R3, and R5 for both suspended sludge biomass and IFAS. Metagenomic analysis of this wastewater treatment plant is previously described^4^.

Extracted RNA samples were submitted to the Georgia Institute of Technology’s Molecular Evolution Core for rRNA depletion, and RNA sequencing. RNA integrity, read length, and quantification were performed on a 2100 Bioanalyzer with C0.1.069 firmware (Agilent, Santa Clara, California, USA). Excess rRNA was depleted using QiaSeq FastSelect 5S/16S/23S depletion kit as well as the FastSelect HMR and Plant rRNA probes (Qiagen, Hilden, Germany). RNA sequencing was performed on an Illumina NovaSeq S1 PE100bp to obtain 1.6 billion reads (Illumina Inc, San Diego, California, USA). Samples were demultiplex and barcodes removed by the core facilities prior to releasing sequences.

Raw reads were preprocessed using fastp v0.23.4^43^ before residual rRNA reads were separated from mRNA reads using SortMeRNA v2.1^44^. The remaining mRNA reads were then competitively mapped to previously assembled metagenome-assembled genomes (MAGs) from AOB, CMX, NOB, and AMX^4,14,18^. The MAGs were annotated using bakta v1.8.2^45^. The mapped mRNA reads were converted into .bam files, sorted, and indexed using samtools v1.18^46^. Finally, the mapped mRNA reads were quantified for read counts on the annotated MAGs using dirseq v0.43^47^.

### 2.7 Statistical Methods

Statistical analysis was performed in RStudio^30^ with the vegan package^48^. ANOVA was used to determine the impact of DO and reactor zonation across all conditions, while Tukey Honest Significant Differences^49^ post-hoc test was used to compare means between two subgroups. For correlative regression analysis, ranked-based linear regressions were performed in R to determine significant trends across RT-qPCR samples. Shannon Diversity Index^50^ was additionally used to compare the alpha diversity across all amplicon sequencing samples. Differential gene expression analysis was performed with Deseq2^51^. For CMX, the minimum read counts per gene were set to five due to lower transcriptomic reads, whereas the default of 10 read counts was used for AMX.

### 2.8 Data Availability

All code for amplicon sequencing, metatranscriptomic analysis, statistical analysis and plots are available at github.com/queermsfrizzle/CMX_AMX_transcriptomics. Raw fastq files for metatranscriptomic sequencing and amplicon sequencing are available via NCBI bioproject submission number PRJNA1050761.

## 3- Results and Discussion

### 3.1 Microbial community composition and activity was associated with growth phase and DO concentrations

The total and active microbial community composition was distinct from each other between the two phases (i.e., suspended sludge biomass vs attached IFAS phase) based on 16S rRNA gene and transcript composition (**Figure 1**). The largest difference in community composition was between IFAS and suspended sludge biomass which (**Figure 1a**) and explained 27.6% of the total variance between samples (PERMANOVA p << 0.05). There was also a significant difference between each community’s16S rRNA transcript composition and 16S rRNA gene composition which explained 9.6% of the total variance (**Figure 1a**) (PERMANOVA p << 0.05). This is consistent with previous reports which showed the most significant changes to community composition based on reactor zonation followed by the separation of DNA and RNA, and further by the experimental variable of seasonality^38^. When analyzing the Shannon Diversity Index of 16S rRNA transcripts and genes, there was a significantly higher amount of diversity in 16S rRNA genes, compared to transcripts (ANOVA p << 0.05) (**Supplemental Figure 4**). This suggests that a large proportion of the microbial community is likely less active.^52^

**Figure 1:**
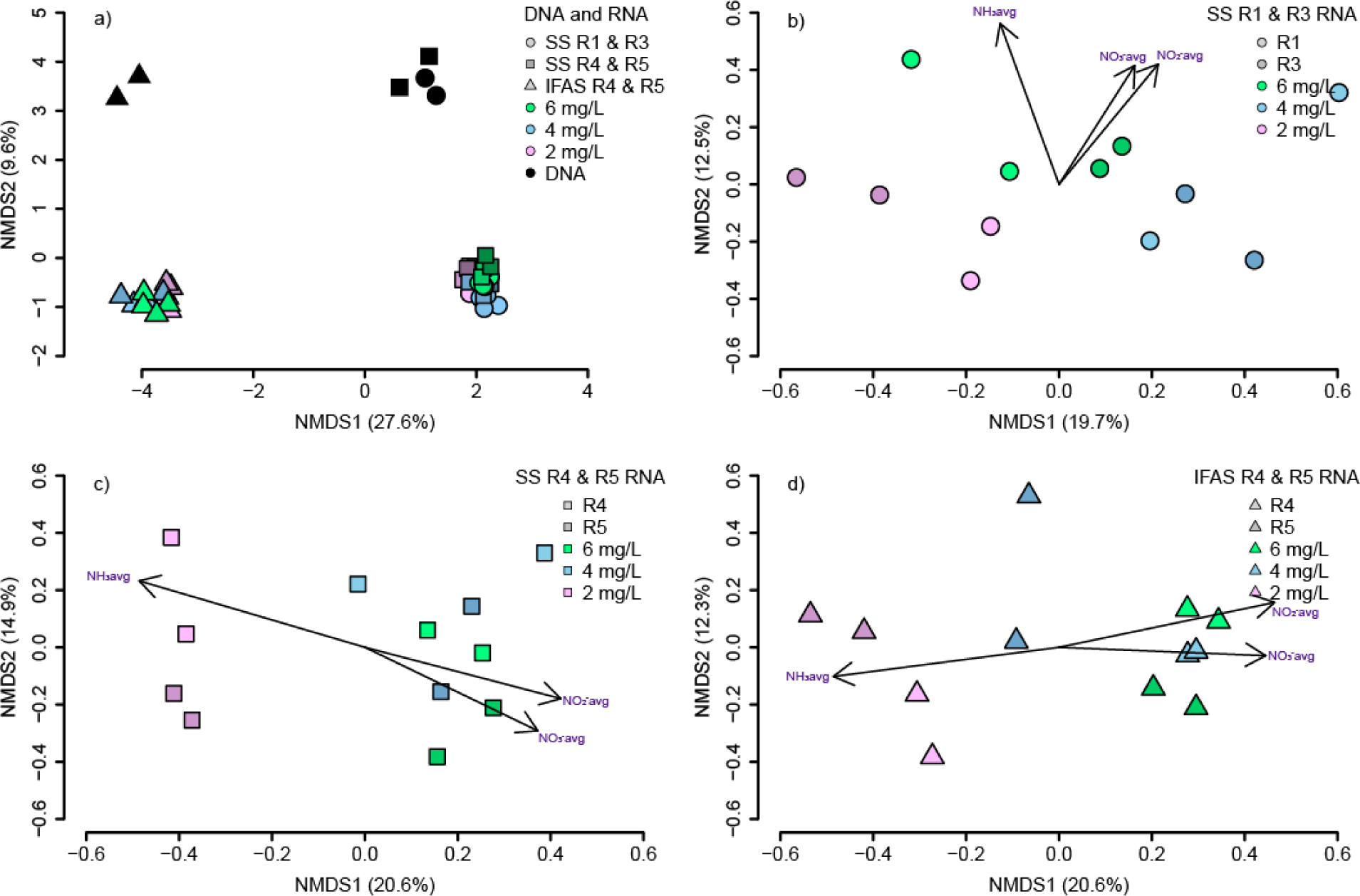
NMDS plots constructed using Bray-Curtis dissimilarity estimates for a) all 16S rRNA gene and transcript OTUs, b) SS for R1 and R3 16S rRNA transcript OTUs, c) SS for R4 and R5 16S rRNA transcript OTUs, and d) IFAS R4 and R5 16S rRNA transcript OTUs. Black data points represent genes while colored data points represent transcripts. Green, blue, and pink shades represent the dissolved oxygen concentration at 6 mg/L, 4 mg/L and 2 mg/L, respectively. Shades of each color represent different specific reactor locations while circles, squares and triangles represent the reactor zones.

For both IFAS biofilms and suspended sludge biomass communities, the DO concentration had a significant impact on the active community composition (PERMANOVA p_IFAS_ = 0.0015, p_ss_ = 0.0007) (**Figure 1, c, & d**). For the IFAS biofilm community, there was a statistically significant difference between DO = 2 mg/L with DO = 4 mg/L as well as DO = 6 mg/L (ANOSIM p_IFASD2D4_ = 0.027 and p_IFASD2D6_ = 0.003), but not between DO =4 mg/L and DO = 6 mg/L (ANOSIM p_IFASD4D6_ = 0.5687). In suspended sludge biomass, all dissolved oxygen concentrations showed statistically significant separation (ANOSIM p_SSD2D4_ << 0.05, p_SSD2D6_ << 0.05, and p_SSD4D6_ = 0.0127). While the DO concentrations did result in significant changes, the measured concentrations of DO, ammonia, nitrate, nitrite, alkalinity, and COD all showed statistically significant effects (all Mantels test p << 0.05) on differences between the active community composition between samples from the IFAS media or suspended sludge biomass. This indicates that the DO setpoint has cascading effects on microbial community activity and redox conditions due to residual oxygen, nitrite, and nitrate. Similar impacts of dissolved oxygen impacting the microbial community composition has been previously shown.^53–55^

### 3.2 High diversity of nitrite oxidizing bacteria compared to strict AOB and AMX

The diversity of nitrifying bacteria was analyzed using a combination of 16S rRNA transcript sequencing and quantification and amplicon sequencing of major nitrogen cycling functional genes (**Figure 2**). 16S rRNA transcript analysis averaged 63,049 ± 11,365 postprocessed reads per sample (**Supplemental Figure 3**) and revealed one dominant AMX (**Figure 2a),** two dominant AOB (**Figure 2c**) and three *Nitrospira*-like OTUs (**Figure 2b**).

**Figure 2:**
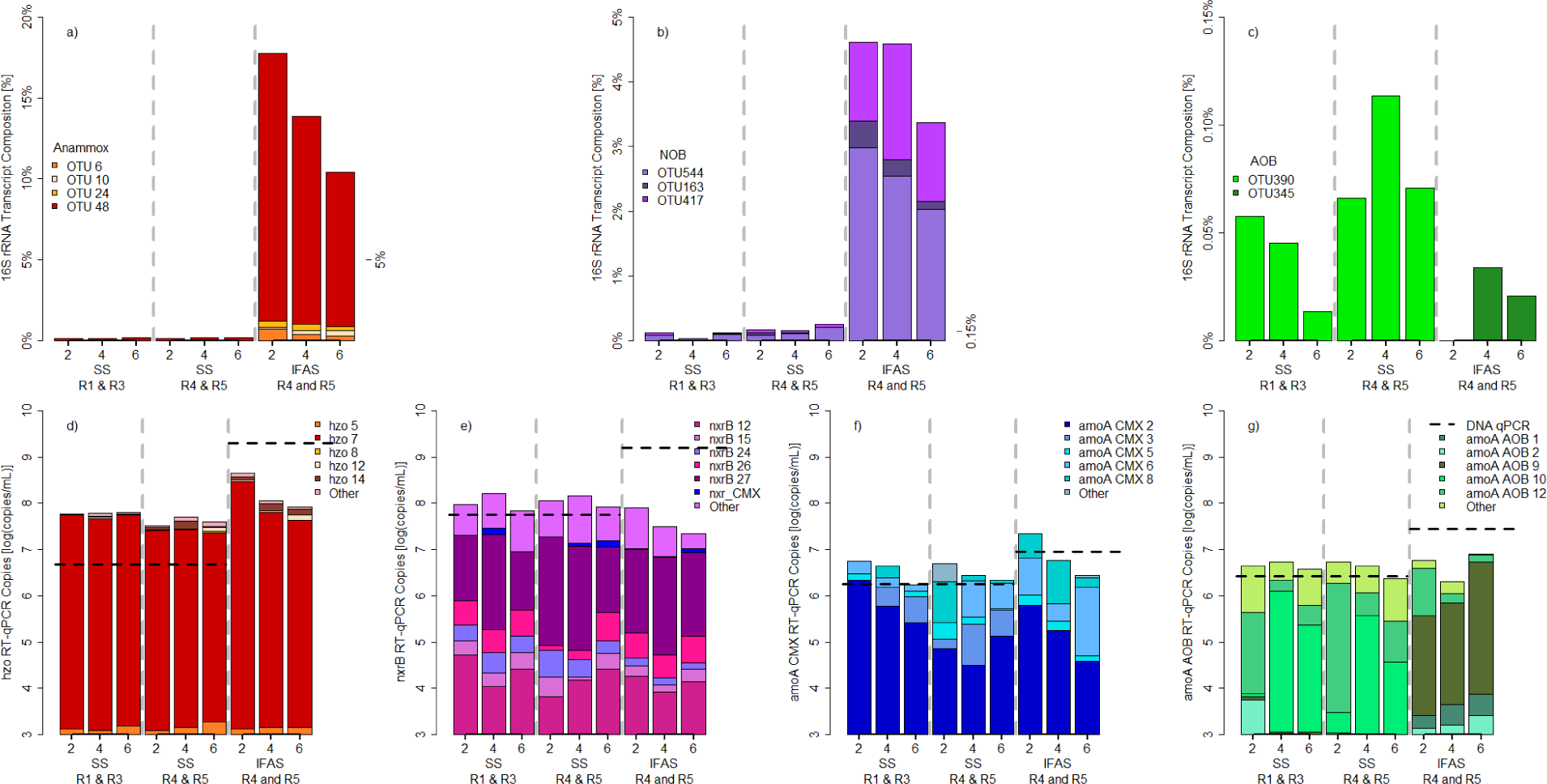
Summary of dominant nitrifying bacteria’s transcriptional activity based on 16S rRNA transcript sequencing (a,b,c) and functional gene expression (d,e,f,g). 16S rRNA transcript activity with a) anammox bacteria b) all nitrite oxidizing bacteria (including comammox which cannot be differentiated using OTUs of 16S rRNA V4 amplicons), and c) strict ammonia oxidizing bacteria. The expression of functional genes are quantified via RT-qPCR and then multiplied with OTU relative abundance in amplicon sequencing data (y-axis) for d) hydrazine oxidoreductase found in AMX e) nitrite oxidoreducatase of all nitrite oxidizing bacteria (including comammox *Nitrospira*), f) ammonia monooxygenase found in comammox *Nitrospira* and g) ammonia monooxygenase found in strict ammonia oxidizing bacteria. The dashed line represents the average abundance of each functional gene via qPCR to determine transcripts-to-gene ratios.

Anammox activity was dominated by OTU48, *Candidatus* Brocadia, based on 16S rRNA transcript quantification. OTU48 was particularly active in IFAS and comprised 16.5% ± 4.2% of the 16S rRNA transcript composition at low dissolved oxygen and declined down to 9.5% ± 2.2% at higher dissolved oxygen concentrations. Similarly, there was a single OTU, *hzo* 7, which dominated anammox activity based on *hzo* transcript absolute abundance (**Figure 2d**) and consistently comprised 93.43% ± 2.85% of the *hzo* transcript composition. IFAS had significantly higher *hzo* expression at low dissolved oxygen (ANOVA p << 0.05) at 8.64 ± 0.13 log(copies/mL) and decreased with higher dissolved oxygen to 7.92 ± 0.13 log(copies/mL). AMX’s *hzo* expression was stable in the suspended sludge biomass at 7.69 ± 0.24 log(copies/mL) and not impacted by dissolved oxygen concentration (ANOVA p = 0.337). While AMX preferring lower dissolved oxygen concentrations has been previously reported^23,56^, what is interesting when comparing our system is the lack of diversity in AMX, whereas typically multiple species of abundant AMX^57^ are observed to co-occur from wastewater bioreactors seeing Ca. *Brocadia* and Ca. *Kuenenia* together^56^, to rice-wheat paddy fields observing Ca. *Brocadia* and Ca. *Jettenia*^58^. While there are low abundance species observed (**Supplemental Figure 4**), these species are orders of magnitude lower in abundance and also lower levels of activity.

Three dominant *Nitrospira* OTUs were observed based on 16S rRNA transcript analysis: OTU544 closely associated with with *Nitrospira* defluvii, OTU163 was associated with *Nitrospira moscoviensis*, and OTU417 was unknown *Nitrospira* (**Figure 2b**). For redundancy, 16S rRNA sequences were compared across SILVA SSU 138.1 and MIDAS 5.0 databases to assess whether CMX associated 16S rRNA sequences were distinguishable from NOB, but this was not feasible (data not shown). All 16S rRNA transcripts for *Nitrospira* were significantly more active in IFAS biofilm than in suspended sludge biomass (ANOVA p << 0.05).

In contrast to the 16S rRNA transcript data which showed remarkable differences in *Nitrospira* activity in IFAS biofilm as compared to suspended sludge biomass, RT-qPCR data indicates relative stable transcriptional activity of the *nxrB* gene in the suspended sludge biomass (7.99 ± 0.23 log(copies/mL)) which was comparable to that of the IFAS biofilm based *nxrB* activity (**Figure 2e**). Interestingly, *nxrB* transcript abundance dropped significantly with increasing dissolved oxygen (ANOVA p << 0.05) from 7.89 ± 0.08 log(copies/mL) to 7.31 ± 0.12 log(copies/mL) from DO = 2 mg/L up to DO = 6 mg/L in the IFAS biofilm community. If *nxrB* activity was only associated with nitrite oxidation, this drop would be perplexing since ammonia oxidation is higher at high dissolved oxygen concentrations. However, this drop in *nxrB* transcripts could be indicative of lower nitrite concentrations found in IFAS biofilm since the available nitrite is likely consumed by AMX, or alternatively in anoxic or low DO conditions *nxrB* could reversibly reduce nitrate back to nitrite. This would explain the higher *nxrB* transcripts in IFAS biofilm at lower dissolved oxygen concentrations.

To determine if any *nxrB* transcripts were from CMX, the *nxrB* transcripts were mapped to all *Nitrospira* MAGS. Several low abundance *nxrB* OTUs mapped to CMX accounting for 1.02% ± 0.84% of the *nxrB* transcript composition without significant changes due to dissolved oxygen concentration (ANOVA p = 0.83). Interestingly, the *amoA* CMX activity was significantly higher in the IFAS biofilm compared to the suspended phase and demonstrated sensitivity to DO concentration (ANOVA p = 0.033) increasing from 6.43 ± 0.32 log(copies/mL) up to 7.33 ± 0.58 log(copies/mL) with drop in DO from 6 to 2 mg/L (**Figure 2f**). Overall, in suspended sludge biomass CMX appear to have a stable transcriptional activity of *nxrB* and a low activity of *amoA* whereas in IFAS biofilm a significant increase in *amoA* activity is observed. Interestingly, CMX *amoA* activity did significantly increase at reduced DO, even in suspended sludge biomass. This is uncommon since *amoA* activity is regulated by ammonia concentration, not oxygen concentration. This change in expression in the suspended phase is likely due to residual *amoA* transcripts of biomass being sloughed off from IFAS biofilm into the suspended phase which would experience fluctuations in ammonia concentration, therefore impacting *amoA* transcript regulation based on DO concentration.

This aligns with current studies showing that CMX strongly prefer biofilms^59^ and our previous work using intrinsic assays and differential inhibition to suggest higher ammonia oxidation from CMX in IFAS biofilm^4^. Other studies using differential inhibitors with CMX enrichment cultures from rotating biological contactors have shown similar changes in CMX activity between biofilm and suspended sludge biomass cultures^60^. While our study suggests a greater in nitrite oxidation activity of CMX in suspended sludge biomass, future studies are required to investigate the mechanisms underpinning CMX differential expression in suspended and attached phases.

Two dominant strict AOB (OTU390, OTU345) within the family *Nitrosomonadaceae* were detected in the 16S rRNA transcript amplicon sequencing library. While both OTUs showed very low levels of activity based on their relative abundance in the 16S rRNA transcript libraries, OTU390 was only active in the suspended phase, while only OTU345 demonstrated low levels of activity in the IFAS biofilms (**Figure 2c**). While the absolute abundance of AOB was significantly higher in IFAS biofilm samples (7.45 ± 0.9 log(copies/mL)) compared to suspended sludge biomass samples (6.42 ± 0.26 log(copies/mL)), their transcriptional activity based on RT-qPCR assay targeting the *amoA* gene indicated no significant change with either reactor zonation or DO concentration (ANOVA_DO_=0.426, ANOVA_zone_= 0.839) averaging at 6.63 ± 0.31 log(copies/mL) (**Figure 2g**). This indicates a significant downregulation of *amoA* transcriptional activity (on a per cell basis) in the IFAS biofilm community. Interestingly, dissolved oxygen or reactor zonation did not impact the transcriptional activity of *amoA* for AOB. This stable activity of *amoA* by AOB is consistent with past activated sludge systems^61–63^ but overall lower than typically reported between 7-8 log(copies/mL). This is consistent with our previous findings that CMX and AMX significantly contribute to nitrogen removal, especially in IFAS biofilm phase^4^.

Multiple *amoA* OTUs were identified in both the suspended sludge biomass and IFAS biofilm phase which is in contrast with the single 16S rRNA gene OTU in each phase. This could likely emerge due to both the presence of multiple *amoA* gene copies within genomes relative to a single 16S rRNA gene per AOB genome^64^, but also could emerge from the differences in the sensitivities of two assays. Specifically, *amoA* gene amplicon sequencing was performed after selectively amplifying this gene it from cDNA, and as a result is likely to detect sequences from more rare populations as compared to when the entire microbial community is profiled using 16S rRNA gene assay. Nonetheless, in alignment with the 16S rRNA transcript data, a single OTU, *amoA 9,* was primarily active in the IFAS biofilm community. This *amoA 9* transcript exhibited sensitivity to DO concentration increasing from 57.2% at 2 mg/L to 74.4% of all AOB *amoA* transcripts at 6 mg/L. Furthermore, the *amoA* 9 OTU sequence exhibited 99.6% sequence similarity with an uncultured *Nitrosomonas clone A12* ^65^. This *Nitrosomonas clone A12* was previously found in a biofilm working alongside CMX and raises the possibility that specific *Nitrosomonas* species are more inclined to co-exist with CMX or potentially both prefer biofilms due to their growth kinetics.

### 3.3 Comammox Nitrospira and anammox bacteria have strong synergistic relationship in the IFAS biofilm phase

CMX and AMX show a strong symbiotic relationship in IFAS biofilm as indicated by significant correlative relationships (**Figure 3a**) and transcripts-per-million gene expression analysis of major nitrogen cycling pathways (**Figure 3b& c**). Using RT-qPCR transcript abundances, *hzo* transcripts were regressed against *amoA* for CMX transcripts in IFAS biofilm (**Figure 3a**) and a significant ranked linear correlation was observed (linear regression R^2^ = 56.1% & p = 0.003) suggesting that AMX *hzo* and CMX *amoA* activities may be potentially co-regulated by changes in DO concentrations. While similar regressions can be performed using *nxrB,* CMX *amoA* and AMX *hzo* are the only genes that exhibited significantly higher transcriptional activity in the IFAS biofilm relative to the suspended phase. This contrasts with expression levels of AOBs *amoA* and total *nxrB* transcripts which were either stable or declined in the IFAS biofilm, respectively. This suggests that aerobic ammonia oxidation by CMX was strongly associated with anaerobic ammonia oxidation by AMX, suggesting that CMX may play a role in supplying nitrite to AMX. This has been suggested since the discovery of CMX coexisted with an AMX population^66,67^ but this study is the first to demonstrate this symbiotic relationship leveraging transcriptomic expression.

**Figure 3:**
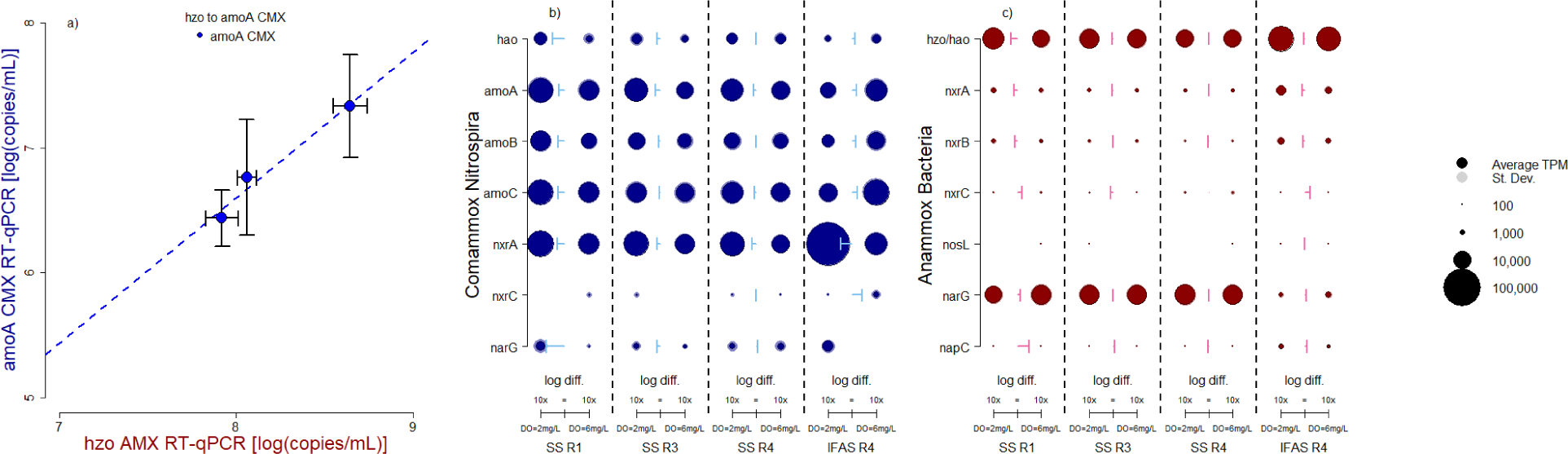
Anammox bacteria and comammox Nitrospira have a strong linear relationship of their activity in IFAS biofilm based on a) RT-qPCR transcript quantification of hydrazine oxidoreductase and ammonia monooxygenase from comammox Nitrospira. This symbiotic relationship was further explored using transcripts-per-million activity profiles of major nitrogen cycling pathways mapped to MAGS constructed of b) comammox Nitrospira and c) anammox bacteria. The size of each point relates to the magnitude of transcripts-per-million, with an outline for standard deviation. The arrows show the magnitude difference between each data point DO = 2 mg/L and DO= 6 mg/L.

This relationship was further investigated by examining differential expression of key nitrogen cycling genes in MAGS from CMX (**Figure 3b**) and AMX (**Figure 3c**) by mapping metatranscriptomic reads to them. Similar analysis for AOB and NOB can be found in the supplemental materials (**Supplemental Figure 6**). Nitrogen cycling genes from CMX were most impacted by changes in DO with significant increases in *nxrA* across every reactor setting (Tukey all p << 0.05) at lower DO concentrations. In suspended sludge biomass, nitrogen cycling pathways accounted for 7.75% ± 1.62% of the CMX transcripts at low dissolved oxygen concentrations and only 4.31% ± 0.85% at high DO concentrations. This was further highlighted when comparing activity in IFAS biofilm with 18.53% ± 2.89% and 7.18% ± 1.40% at DO = 2 mg/L and DO = 6 mg/L of all transcripts respectively mapped to nitrogen cycling pathways. The largest increase in activity was associated with *nxrA* in IFAS biofilm which increased 9.1-fold at low DO concentrations. This was surprising since *nxrC* expression did not significantly change between DO setting and was often undetected but this may be a result of a fragmented MAG resulting in low mapped transcripts since other studies have shown consistent transcriptional activity across all *nxr* subunits^68–70^. Further work needs to investigate biofilm depth-dependent activities and oxygen gradients to understand the how CMX regulates expression of nitrogen cycling genes across a redox gradient.^71^

AMX maintained mostly consistent expression of nitrogen cycling genes in suspended sludge biomass comprising 2.52% ± 0.19% of the total mapped AMX transcripts (**Figure 3c**). In comparison to IFAS biofilm, *narG* (nitrate reductase), was significantly more active in suspended sludge biomass (Tukey p << 0.05) as compared to the IFAS biofilm. This suggests that in suspended sludge biomass AMX may need to supply their own nitrite from residual nitrate due to competition for nitrite with NOB. The downregulation of *narG* in IFAS biofilm could indicate sufficient supply of nitrite, likely from CMX. Additionally, in IFAS biofilm, most pathways experienced a significant increase in transcripts-per-million at low DO concentrations. The expression of genes associated with *hao*/*hzo* pathway increased 1.2x, *nxrA* increased 1.68x, and *nxrB* increased 1.36x. These increases results in AMX’s nitrogen cycling transcripts in IFAS biofilm increasing from 2.65% ± 0.23% at DO = 6 mg/L up to 3.23% ± 0.08% at DO = 2 mg/L (Tukey p << 0.05) with overall activity in IFAS biofilm being significantly higher than in suspended sludge biomass.

### 3.4 Differential Gene Expression Analysis Anammox Increases Flagella Activity in IFAS biofilm phase under high DO conditions

AMX_281 was the most actively transcribed MAG in IFAS biofilm with 93% ± 0.2% of the transcripts mapping at DO = 2 mg/L, and 88.3% ± 3.1% at DO = 6 mg/L (**Figure 4a**). This is consistent with dominance of 16S rRNA transcript OTU 48 whose transcripts completely mapped to the MAG, and the dominance of *hzo* transcript OTU 7, which did not map to the MAG due to an incomplete genome.

**Figure 4:**
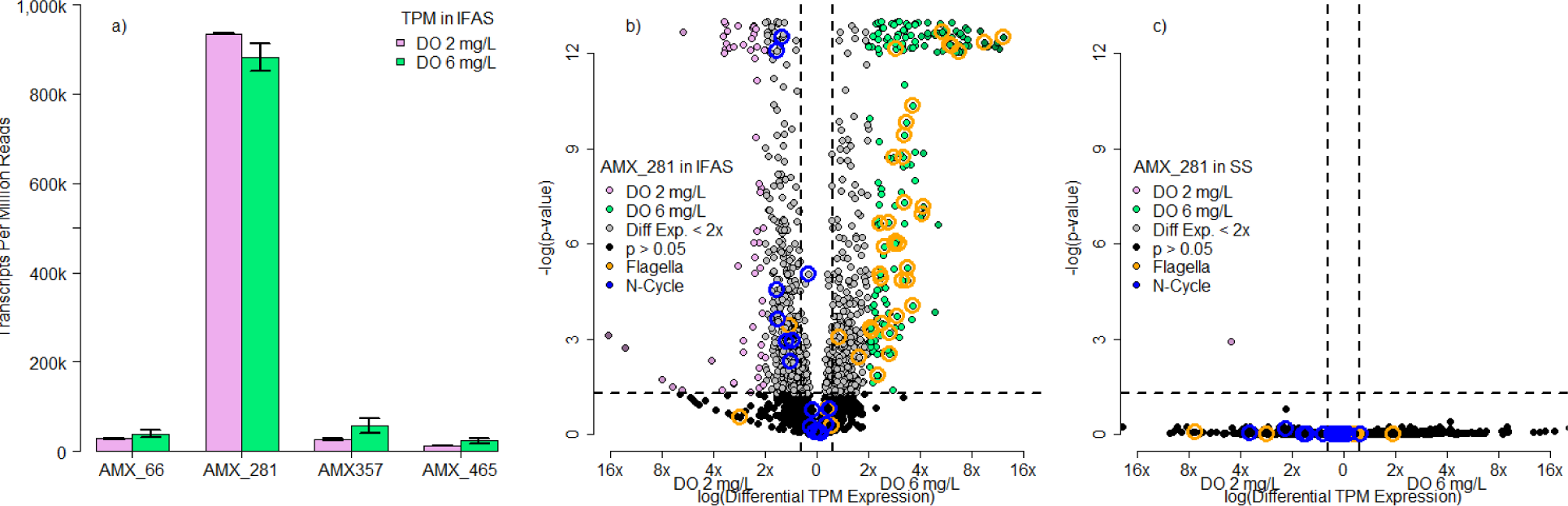
Anammox MAG, AMX_281, was highly expressive in IFAS biofilm compared to other MAGS (Figure 4a), and further analyzed using Deseq2 for differential gene expression analysis in b) IFAS biofilm and c) suspended sludge biomass. Points in pink were statistically significantly more active and expressive at DO = 2 mg/L while points in green were statistically significantly more active and expressive at DO = 6 mg/L. Blue circles represent shifts in nitrate transporting, while the orange circles represent transcripts related to flagella activity. In instances where the -log(p-value) exceeded 12, they are plotted as 12.5 with a jitter for spacing.

AMX_281 was further analyzed in IFAS biofilm (**Figure 4b**) and in suspended sludge biomass (**Figure 4c**) with genes considered significantly differentially transcribed with a Wald test p < 0.05 and greater than 2x transcripts-per-million between two conditions being compared (i.e., 2 vs 6 mg/L of DO or biofilm vs suspended phase). Most genes associated with nitrogen cycling functions in AMX_281 showed insignificant changes in relative expression abundance, similar to all AMX in **Figure 3c**. If nitrogen cycling genes are up-regulated as shown with *hzo* quantification via RT-qPCR in **Figure 2d**, they increase proportionally to maintain similar relative expression abundance via transcripts-per-million and differential gene expression (**Figure 3c** and **Figure 4b**). At lower DO concentrations, most up-regulated gene expression was from hypothetical proteins and was not further investigated.

At high dissolved oxygen concentrations in IFAS biofilm, there was a significant increase in most flagella-related transcriptional activity, which was not similarly observed in suspended sludge biomass. This was observed for genes across multiple flagellar related proteins from FlgN, FliS, Flagellar hook-associated protein 2, and FlaG, all of which increased more than five-fold at high DO concentrations. Yan et al. 2020^23^ suggests this maybe be oxygen related-stress response and trigger biofilm formation. While reactive oxygen species could impact AMX in IFAS biofilm, this stress response is not consistent with AMX in suspended sludge biomass (**Figure 4c**) which did not show significant changes in flagella-related activities and accounted for an overall lower abundance of transcripts-per-million (IFAS_DO6_ = 1.78%, SS_DO6_ = 0.08%). We hypothesize that anammox bacteria in IFAS biofilms may increase mobility due to chemotaxis and cell growth within the competitive environment of a biofilm. Research into anammox located in deep sea anoxic nitrate-ammonium transition zones observed expression of full gene sets for flagella mobility despite limited energy for mobility, suggesting flagellar activity is linked to in situ growth^72^. Similarly, other *Planctomycetes* bacteria have shown increased flagella mobility in highly rich media^73^ suggesting AMX may utilize chemotaxis within high nutrient concentrations as opposed to mobility to avoid oxygen stress^74^.

### 3.5 Comammox Nitrospira alters ammonia and nitrite metabolic preference based on redox conditions

Two CMX MAGS revealed a significant shift in metabolic preferences based on dissolved oxygen concentration (**Figure 5**). CMX_1 was overall more active than CMX_2 comprising 62.5% ± 0.9% of transcripts-per-million at DO = 2 mg/L and 86.4% ± 3.0% at DO = 6 mg/L whereas CMX_2 comprised 37.5% ± 0.9% at DO = 2 mg/L and 13.6% ± 3% at DO= 6 mg/L respectively (**Figure 5a**). Further, genes associated with ammonia oxidation were significantly upregulated at 6 mg/L DO, while genes associated with nitrite oxidation were significantly upregulated at 2 mg/L DO.

**Figure 5:**
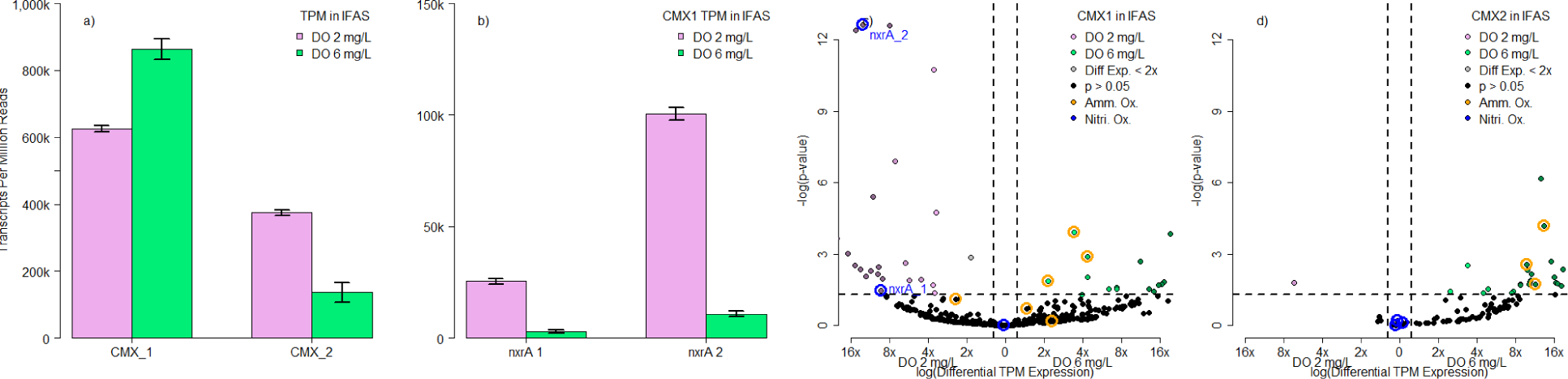
Comammox Nitrospira had two dominant MAGS in IFAS biofilm (Figure 5a) which were further analyzed using volcano plots for differential gene expression analysis in IFAS biofilm. The more abundant MAG, CMX_1, saw significant changes in the expression between the two copies of nxrA within the genome (Figure 5b). Each MAG was analyzed with Deseq2 for differential gene expression analysis in IFAS biofilm with c) CMX_1 and d) CMX_2. Points in pink were statistically significantly more active and expressive at DO = 2 mg/L while points in green were statistically significantly more active and expressive at DO = 6 mg/L. Blue circles represent shifts nitrite oxidation activity, while the orange circles represent shifts in ammonia oxidation activity. In instances where the -log(p-value) exceeded 12, they are plotted as 12.5 with a jitter for spacing.

Both MAGS, CMX_1 in **Figure 5c** and CMX_2 in **Figure 5d**, showed significant increases in transcription of genes involved in nitrite oxidation pathways at lower DO concentrations in IFAS biofilm, further reinforcing *nxrB* increases shown in **Figure 3b**. Interestingly, while this analysis algins with *nxrB* RT-qPCR results (**Figure 2e**), the relative downregulation of *amoA* in CMX gene expression conflicts with *amoA* transcript quantification in CMX via RT-qPCR (**Figure 2f**) which showed an increase in expression. This is likely a result of transcripts-per-million being a relative abundance metric whereas RT-qPCR is quantitative. At low DO levels, CMX increases expression of genes involved with both ammonia oxidation and nitrite oxidation, but those associated with nitrite oxidation have a much larger increase resulting in the expression of genes associated with ammonia oxidation appearing to decrease using relative abundance metrics.

CMX_1 had sufficient mapped transcripts to compare differential expressions between the two copies of *nxrA* within the genome. The second copy of *nxrA* had significantly more activity than the first copy of *nxrA* (Tukey p << 0.05). Furthermore, the ratio of expression between the second copy and the first copy was constant at 4.14x copies and no significant differences in the ratio based on dissolved oxygen concentration (Tukey p = 0.739). This suggests that there are transcription regulations for these two *nxr* pathways. Different regulation for the paralogs has been previously observed in cultures^75^, but not previously reported in a full-scale wastewater treatment system.

Interestingly, the *nxr* pathway is also a reversible pathway, which can either oxidize nitrite or reduce nitrate in anaerobic conditions^76^. This dynamic further reinforces the synergistic relationship shown in **Figure 3** where CMX are likely supplying AMX nitrite in IFAS biofilm as previously reported by Vilardi et al^4^. In aerobic conditions CMX could perform ammonia oxidation and supply nitrite to AMX. In anaerobic conditions, the *nxr* pathway could be reversed for nitrate reduction to resupply nitrite to AMX. This scenario could explain why CMX significantly increases ammonia oxidation and greatly decrease *nxr* expression in IFAS biofilm, whereas strict NOB do not significantly increase *nxrA* expression in IFAS biofilm phase (**Supplemental Figure 6c**). Furthermore, there is an overall decrease in *nxrB* expression in IFAS biofilm compared to suspended sludge biomass (**Figure 2e**) since CMX dominate the limited *Nitrospira* community in IFAS biofilm and strict NOB dominate the suspended sludge biomass. While others have reported differential expression of *amo* and *nxr* in CMX^68,69^, this would be the first report suggesting DO role and redox zonation promoting differential expression.

### 3.6 Implications for development of a CMX-AMX system for mainstream nitrogen removal and future research

The cooperation of CMX and AMX in IFAS biofilms at lower DO concentrations has significant beneficial implications for wastewater treatment facilities. Such coupling effectively doubles as both cost saving due to lower aeration energy consumption^77^, and decarbonization by potentially minimizing the release of nitrous oxide during nitrification^5^. Further, integrating an IFAS biofilm component within conventional nitrification-denitrification process does not require significant infrastructure changes and can be accomplished by integrating carrier media within existing reactors. The metabolic flexibility of CMX serves three critical roles benefiting AMX by 1) aerobically converting ammonia to nitrite for AMX’s anaerobic ammonia oxidation 2) potentially anaerobically converting nitrate back to nitrite for further AMX anaerobic ammonia oxidation and 3) utilizing oxygen via aerobic ammonia oxidation and nitrite oxidation to create anaerobic zones for AMX. While this study demonstrates that nitrifying activities of CMX and their co-operation with AMX are sensitive to DO concentrations, it also opens up further research questions and needs for process optimization. What is the optimal DO concentration to maximize anaerobic ammonia removal considering the more pronounced upregulation of genes associated with nitrite oxidation as compared to ammonia oxidation for CMX bacteria at lower DO concentrations? Specifically, do CMX bacteria start competing with AMX for nitrite below a certain DO setpoint despite substantially lower nitrite affinities^4^. Further, the conditions critical for establishment and maintenance of AMX and CMX co-operation in a full-scale mainstream system are as yet unknown since our work was conducted on already established nitrifying community.

## Acknowledgements

Johnston is supported by the National Science Foundation under Grant # EEC-2127509 to the American Society for Engineering Education. This research is supported by NSF CBET 1703089 and NSF CBET 1923124

