## Supplemental Materials for "Metatranscriptomic analysis reveals synergistic activities of comammox and anammox bacteria in full-scale attached growth nitrogen removal system"

| Target | Name | Length [bp] | Forward [5'-3'] | Reverse [5'-3'] | Citation |
|---|---|---|---|---|---|
| 16S rRNA V4 | 515F / 806R | 292 | GTGYCAGCMGCC GCGGTAA | GGACTACNVGGG TWTCTAAT | Walters 2015[25] |
| amoA AOB | amoA-1Fmod / amoA_2R | 491 | CTGGGGTTTCTAC TGGTGGTC | CCCCTCKGSAAA GCCTTCTTC | Meinhardt 2015[26] |
| amoA CMX | 496F / 812R | 345 | GCGATTCTGTTTT ATCCCAGCAAC | CCGTGTGCTAAC GTGGCG | Beach 2019[27] |
| nxrB | nxrB169f / nxrB638r | 485 | TACATGTGGTGGA ACA | CGGTTCTGGTCR ATC | Pester 2013[28] |
| hzo | hzocl1F1 / hzocl1F2 | 471 | TGYAAGACYTGY CAYTGG | ACTCCAGATRTG CTGACC | Yang 2020[29] |

**Supplemental Table 1 Primer sequences and amplicon lengths used for qPCR, RT-qPCR, and amplicon gene sequencing analysis.**

| OTU | Blast Description | Sequence ID | Percent Identity |
|---|---|---|---|
| **amoA AOB 1** | Uncultured Clone BB405 | KJ562489.1 | 99.39% |
| **amoA AOB 2** | Uncultured Clone QH2 55 | KT023767.1 | 99.59% |
| **amoA AOB 9** | Nitrosomonas sp. Clone A12_beta | KX024803.1 | 99.39% |
| **amoA AOB 10** | Uncultured Clone amoA_SBR_JJY_11 | FJ577851.1 | 99.80% |
| **amoA AOB 12** | Nitrosococcus sp. Clone SBA21 | KC769050.1 | 99.59% |
| **amoA CMX 2** | Uncultured Nitrospira sp. Clone ZSCMX | ON551360.1 | 98.84% |
| **amoA CMX 3** | Uncultured Nitrospira sp. Clone ZSCMX | ON551360.1 | 98.26% |
| **amoA CMX 5** | Uncultured Nitrospira sp. Clone ZSCMX | ON551360.1 | 99.71% |
| **amoA CMX 6** | Uncultured Nitrospira isolate RBC Group J | LR723642.1 | 98.55% |
| **amoA CMX 8** | Uncultured Nitrospira isolate RBC Group J | LR723642.1 | 97.39% |
| **nxrB NOB 12** | Uncultured Nitrospira sp. Clone HKA-E8 | KC884878.1 | 99.58% |
| **nxrB NOB 15** | Uncultured bacterium nsxB-31 | AB846874.1 | 99.58% |
| **nxrB NOB 24** | MAG: Nitrospira sp. Isolate UBC3 chromosome | CP092079.1 | 97.32% |
| **nxrB NOB 26** | Uncultured bacterium nsxB-04 | AB846849.1 | 99.17% |
| **nxrB NOB 27** | Uncultured bacterium NOB OTU149 NLJ4_2973 | MW343074.1 | 98.68% |
| **nxrB CMX 17** | Uncultured prokaryote nxrB | LC257062.1 | 98.01% |
| **hzo AMX 5** | Uncultured planctomycete ANAHZO-4 | EU294367.1 | 98.94% |
| **hzo AMX 7** | MAG: Planctomycetia isolate H1_AMX1 | CP054188.1 | 99.57% |
| **hzo AMX 8** | MAG: Planctomycetia isolate H1_AMX1 | CP054188.1 | 95.10% |
| **hzo AMX 12** | Candidatus Brocadia pituitae complete genome | AP021856.1 | 99.15% |
| **hzo AMX 14** | MAG: Brocadia sp. Ega_18_Q3-R5-49_MAXAC.112v2 | CP064969.1 | 98.72% |

**Supplemental Table 2 The top BLAST query, sorted based on lowest E-value for representative sequences from each OTU cluster. The matching sequence ID listed with the percent identity similarity.**

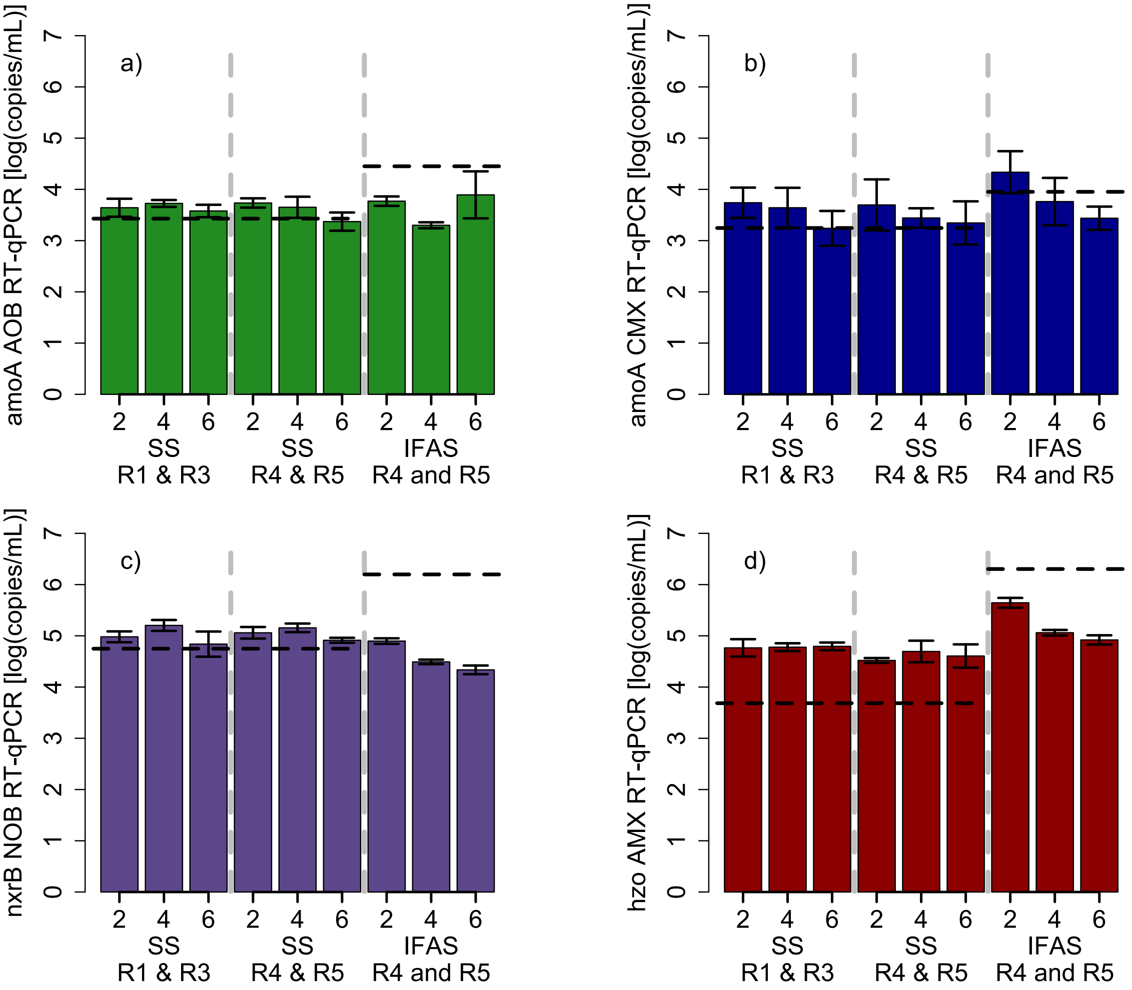

**Supplemental Figure 1 Quantification of functional gene expression via RT-qPCR for a) *amoA* in strict ammonia oxidizing bacteria b) *amoA* in Commamox Nitrospira c) *nxrB* in all nitritie oxidizing bacteria, and d) *hzo* in anammox bacteria. SS = suspended sludge, IFAS = IFAS biofilms. The standard deviations are shown with the error bars and the average functional genes abundance via qPCR is shown with the dashed lines.**

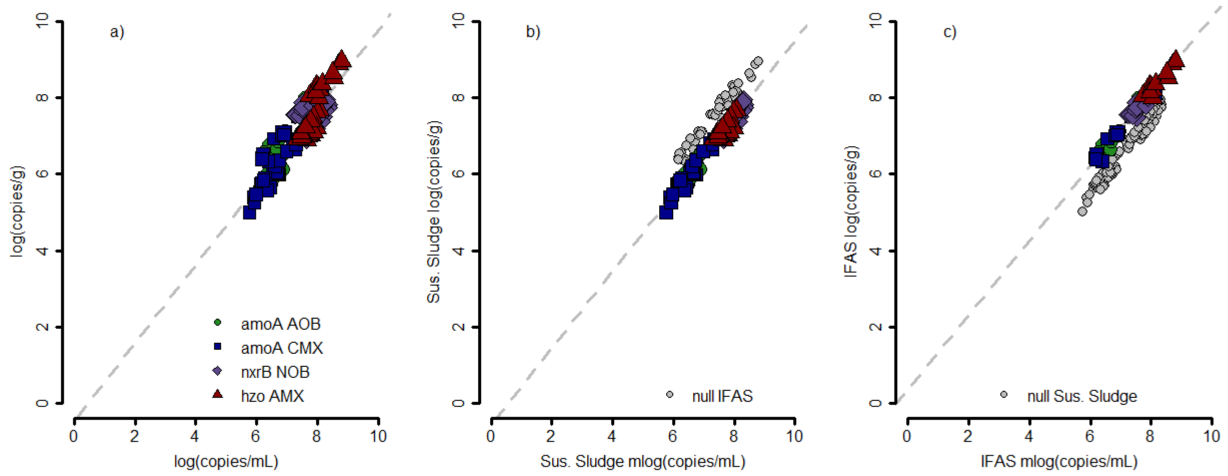

**Supplemental Figure S2 Comparison of normalization techniques between using copies per volume vs copies per mass of sludge. Suspended sludge b) has a strong linear relationship with $R^2 = 0.963$ and $p << 0.05$. The average ratio of log(copies/gram)/log(copies/mL) was 0.927 ± 0.022 for suspended sludge. In IFAS c), the strong linear relationship was also observed with an $R^2 = 0.972$ and $p << 0.05$. The average ratio of log(copies/gram)/log(copies/mL) was higher 1.024 ± 0.018 because the biomass on IFAS was significantly more dense than the suspended sludge.**

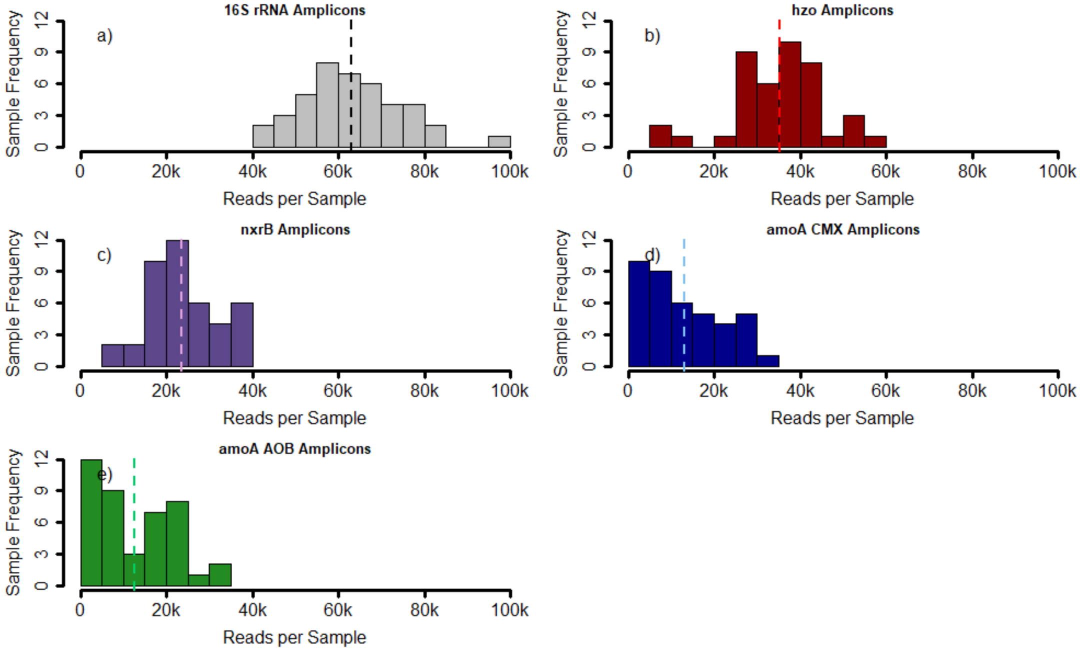

**Supplemental Figure 3 The frequency and distribution of post-processed and filtered reads a) 16S rRNA transcript sequences, b) *hzo* transcript sequences, c) *nxrB* transcript sequences, d) *amoA* for Comammox *Nitrospira* transcript sequences and e) *amoA* for strict ammonia oxidizing bacteria transcript sequences. Dashed lines indicate the average reads per sample.**

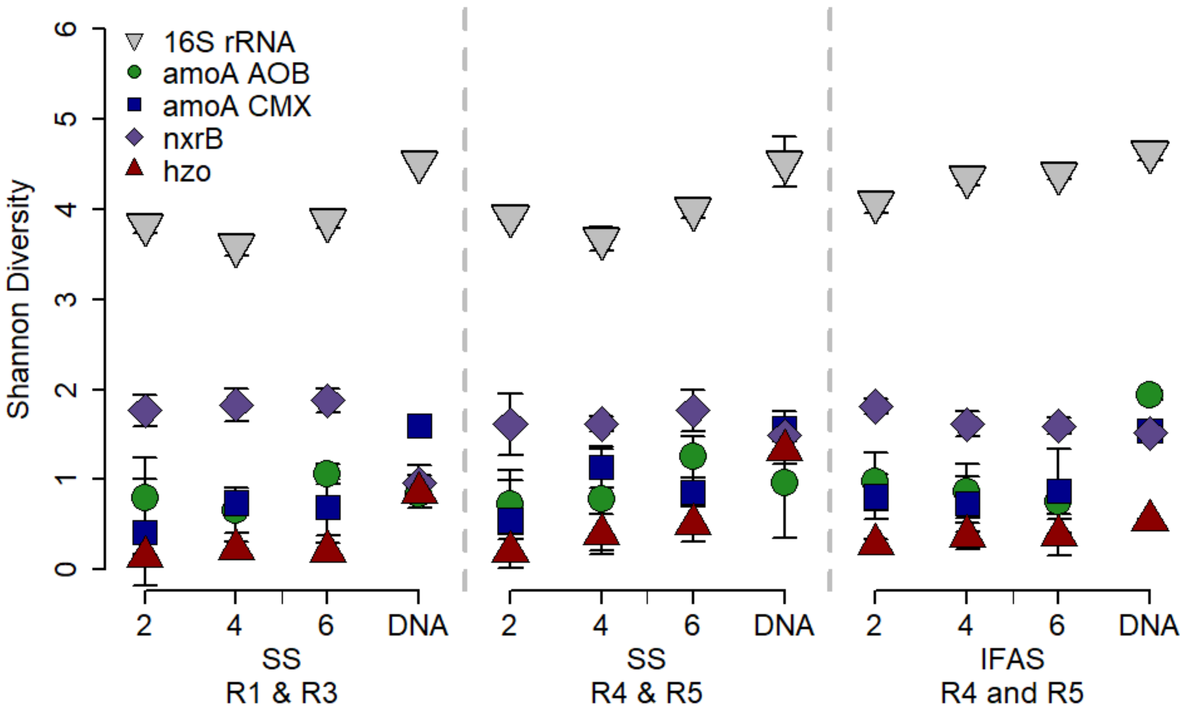

**Supplemental Figure 4 Shannon diversity indexes for the compositional analysis of 16S rRNA in gray inverted trianges, *amoA* for ammonia oxidizing bacteria in green circles, *amoA* for Comammox *Nitrospira* in blue squares, *nxrB* for nitrite oxidizing bacteria in purple diamonds, and *hzo* for anammox bacteria in red triangles.**

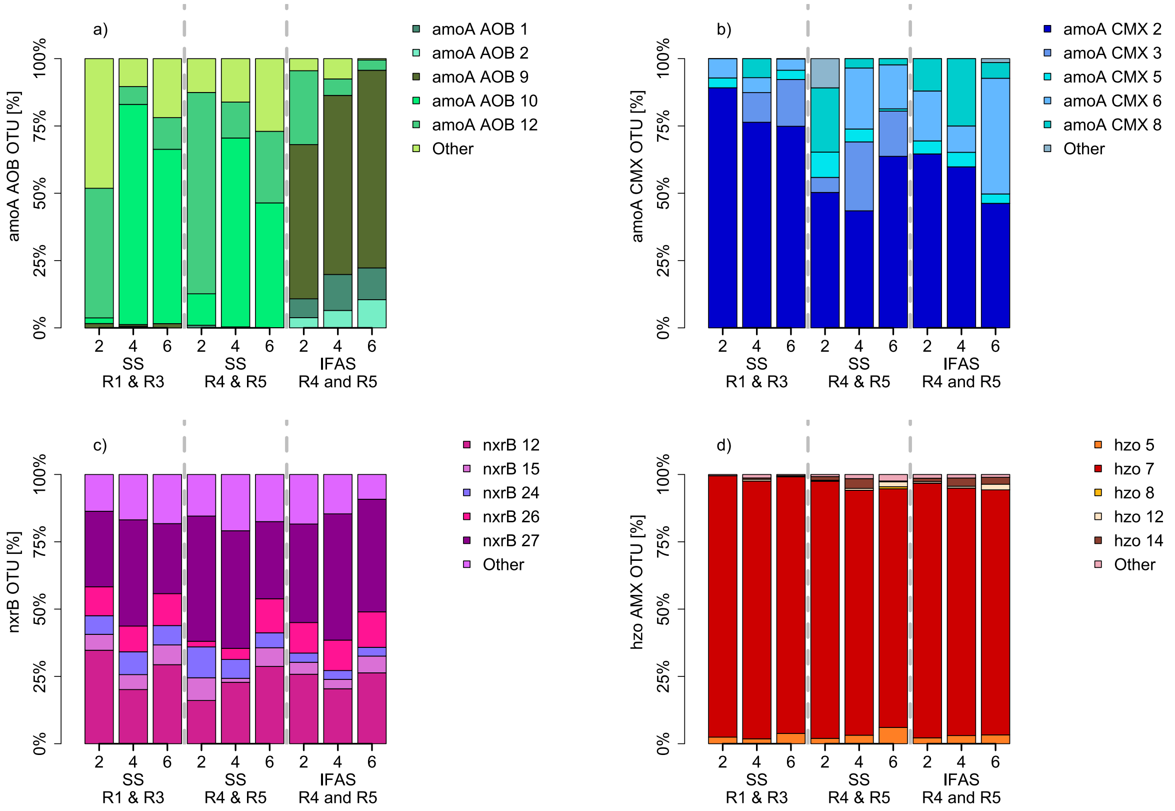

**Supplemental Figure 5. Composition of functional gene expression via amplicon transcript sequencing for a) *amoA* in strict ammonia oxidizing bacteria b) *amoA* in Commamox *Nitrospira* c) *nxrB* in all nitritie oxidizing bacteria, and d) *hzo* in anammox bacteria.**

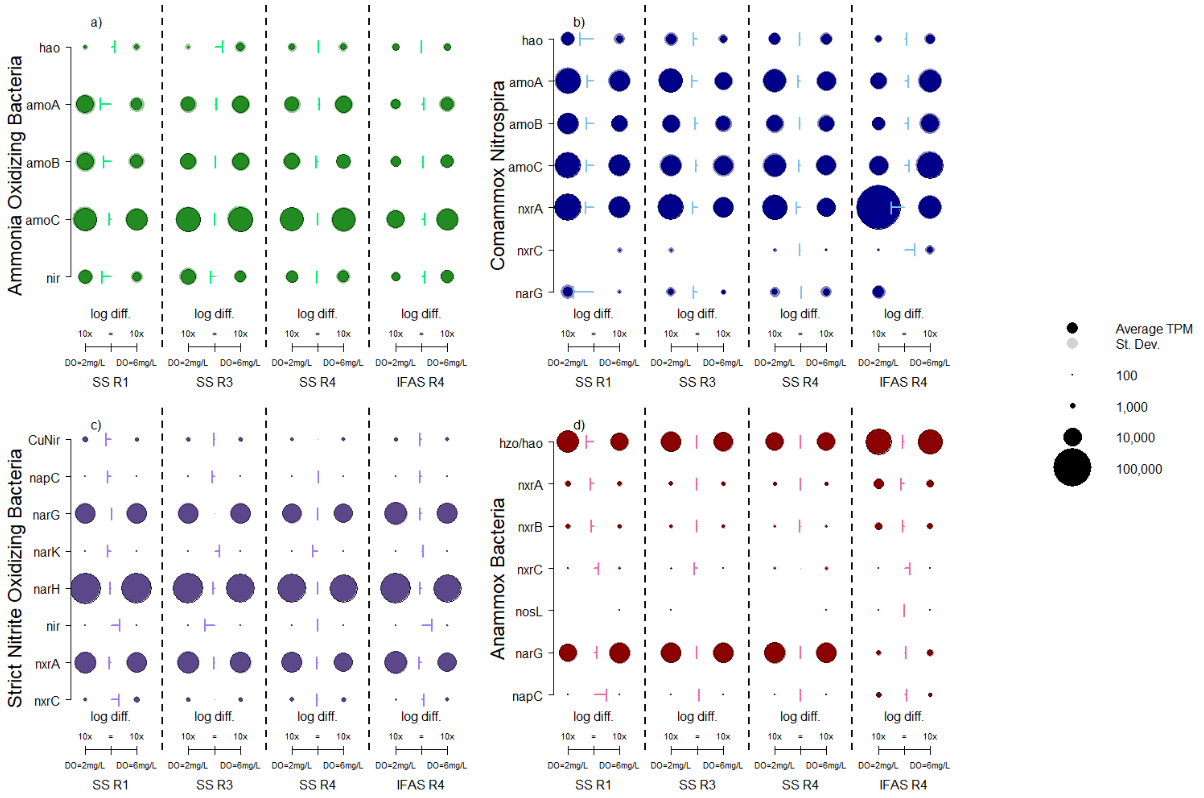

**Supplemental Figure 6 Transcripts-per-million activity profiles of major nitrogen cycling pathways mapped to MAGS constructed of a) strict ammonia oxidizing bacteria, b) comammox *Nitrospira,* c) strict nitrite oxidizing bacteria and d) anammox bacteria. The size of each point relates to the magnitude of transcripts-per-million, with an outline for standard deviation. The arrows show the magnitude difference between each data points DO = 2 mg/L and DO= 6 mg/L.**

*Strict ammonia oxidizing bacteria and strict nitrite oxidizing bacteria dominate suspended sludge activity*

Transcripts were mapped to previously assembled MAGS to determine the impacts of dissolved oxygen concentration on major nitrogen cycling pathways (**Supplemental Figure 6**). The nitrogen cycling pathways of ammonia oxidizing bacteria in suspended sludge were mostly unimpacted by dissolved oxygen concentration with nitrogen cycling transcripts comprising 3.76% ± 0.82% of all mapped AOB transcripts (**Supplemental Figure 6a**). The only exception was with *nir,* nitrite reductase, which significantly increased at low dissolved oxygen in R1 (Tukey p = 0.014). In IFAS at high dissolved oxygen concentrations, *amoC* and *nir* were significantly upregulated (Tukey $p_{amoC} = 0.002$, $p_{nir} = 0.001$). Overall nitrogen cycling expression in ammonia oxidizing bacteria did decrease in IFAS down to 1.77% ± 0.28% at DO = 2 mg/L and 3.04% ± 0.23% at DI = 6 mg/L.

Strict NOB were significantly more active at high dissolved oxygen concentrations in all systems (Tukey all p << 0.05) (**Supplemental Figure 6c**). The percentage of nitrogen cycling transcripts mapped to the MAGS comprised 6.87% ± 0.63% in suspended sludge and 7.64% ± 0.68% in IFAS at DO = 2 mg/L, down to 6.24% ± 0.67% in suspended sludge and 5.70 ± 0.09% in IFAS at DO = 6 mg/L. Primarily, *nxrA,* and *narH* were most upregulated at low dissolved oxygen concentrations suggesting incomplete nitrogen removal causing an accumulation of nitrate and nitrite. While these two pathways did significantly increase at low dissolved oxygen concentrations, their largest increases were only 1.52x and 1.33x in IFAS respectively.
